# Amyloid-beta deposition and reduced drainage at the cribriform plate lymphatics in APP/PS1 mouse model of Alzheimer’s Disease

**DOI:** 10.1101/2025.05.15.654007

**Authors:** Sophia M. Vrba, Collin Laaker, Ajinkya R. Limkar, Martin Hsu, William A. Ricke, Tyler K. Ulland, Mariana Pehar, Matyas Sandor, Zsuzsanna Fabry

## Abstract

Alzheimer’s disease (AD) is the most common cause of dementia, leading to substantial personal, economic, and medical costs to patients and society; it is characterized by the build-up of toxic amyloid-beta (Aβ) and hyperphosphorylated tau. It is crucial to the health of the brain that these proteins are processed or drained effectively, but mounting research has shown that in AD pathology there is dysfunction in the ability of the brain to effectively clear pathological Aβ and tau. In this report, we detail the involvement of one important brain drainage pathway and potential site of Aβ clearance, the cribriform plate lymphatics, in 24-month old APP/PS1 mice. We show that cerebrospinal fluid (CSF) efflux is decreased across the cribriform plate area utilizing multiple methods. Moreover, we demonstrate that Aβ aggregates at the cribriform plate – coating surface of olfactory bulbs (OB), olfactory nerve (ON) bundles, and cribriform plate lymphatic endothelial cells (cpLECs). At 24-months, APP/PS1 mice have increased CD45+ cell infiltration and decreased LYVE-1+ vessel area at the cribriform plate, suggesting local inflammation and lymphatic atrophy. Additionally, cpLECs have higher expression of caspase-3 suggesting the decreased LYVE-1 area is due to cellular toxicity resulting in apoptosis. This study demonstrates that the cribriform plate is an important area for further research elucidating its contribution to AD disease pathogenesis.

## Introduction

Alzheimer’s disease (AD) is the most common form of dementia, affecting 60 million people worldwide, with 7 million living in the United States^1^. It is estimated that prevalence will double by 2060 exacerbating the current public health crisis^2^. There are currently no curative treatments for AD, and current treatment options have limited impact on disease progression. Acetylcholinesterase inhibitors (AChEIs)^3, 4^ are mainly symptomatic treatments, and when used in combination N-methyl-D-aspartate receptor antagonists, 5-year survival probability increased by 6%^5^. Recently developed monoclonal antibodies (mAB) against amyloid-beta (Aβ), lecanemab and donanemab, have moderate reduction in cognitive decline^6^, with lecanemab resulting in 25% slower decline in cognitive function^7^. However, there are concerns regarding side-effects with mAB treatments, particularly with amyloid-related imaging abnormalities (ARIA) with 6.1% of donanemab patients experiencing symptomatic ARIA with cerebral edema (ARIA-E)^6^ and 17.3% of lecanemab patients experiencing ARIA cerebral microhemorrhages or superficial siderosis^7^. Therefore, there is an urgent need to better understand AD disease pathogenesis to develop novel therapeutics.

Since Alois Alzheimer’s initial description of Alzheimer’s disease^8^, extensive research into AD pathogenesis has led to the development of several theories on disease progression, and it is widely accepted that deposition of Aβ and hyperphosphorylated tau are pathological hallmarks of the disease leading to cerebral cortical atrophy and loss of neurons. The current dominant theory in the field, the amyloid hypothesis, proposes that AD patients will have an anomalous accumulation of Aβ^9–11^. The abnormal accumulation of Aβ is driven by 1) familial mutations in amyloid precursor protein (APP) or in presenilin 1 or 2 which are involved in APP processing^12–14^ or 2) failure to clear Aβ with ApoE4 and faulty Aβ degradation^15^. Patients with familial mutations present with dementia symptoms in their 40s and 50s, which is termed early-onset AD (EOAD) or familial AD^12–14^. The vast majority of patients, over 90%^16^, present after age 65 and this is termed late-onset Alzheimer’s disease (LOAD) or sporadic AD. LOAD is a complex and multifactorial disease with an estimated heritability of 60-80%^17^ and has been linked to mutations in genes involved in cholesterol metabolism, immune response regulation, and endocytosis^11^. APOE ε4 has been identified as the strongest genetic risk factor for LOAD^18–21^, and its role is not completely understood, but APOE has been shown to participate in removal of Aβ peptides and the ε4 variant is more rapidly catabolized resulting in less APOE to bind Aβ^22^. Overall, despite differences in the underlying genetics driving AD, both LOAD and EOAD result in an increase in Aβ^15^. It is thought that extraneuronal deposition of Aβ can trigger phosphorylation of tau^23^.

Tau, a protein that is involved in regulating microtubule stability in neurons, becomes hyperphosphorylated resulting in microtubule instability, tau misfolding, and aggregation into neurofibrillary tangles^24^. These aggregated forms of Aβ and tau are toxic to neurons^25,26^. Concurrent with the neurodegeneration, the deposition of Aβ and tau are very inflammatory resulting in microglial and astrocyte activation^26–30^. Overall, this pathway leads to neuronal loss and cortical degeneration beginning in the hippocampus and entorhinal cortex, areas involved in memory, and can progress to affect the cerebral cortex, parietal lobe, and temporal lobe.

The failure to clear Aβ and its accumulation is a canon event in disease pathogenesis. While clearance of Aβ is multifactorial, research has focused on the potential of modulating cerebrospinal fluid (CSF) efflux pathways, namely the central nervous system (CNS)-associated lymphatics, to improve Aβ drainage with the intent of slowing disease progression. The field of CNS-associated lymphatics has been reinvigorated with research detailing their importance as a major drainage and reabsorption site of CSF, as well as site of antigen sampling to inform immune responses^31–35^. While the brain parenchyma lacks traditional lymphatic vessels, several CNS-associated lymphatic drainage systems have been mapped out. Drainage systems that access the CSF can be divided into the meningeal lymphatics, which samples through arachnoid cuffs at bridging veins^32,36,37^; the glymphatic system, which relies upon astrocytes, through regulation of aquaporin-4 channels, leading to fluid resorption into perivascular spaces^38,39^; and cranial nerve and perineural lymphatic pathways, which have shown CSF exit in the perineural spaces along cranial nerves^31,35,40–42^. The meningeal lymphatics, which exist in the dural layer above the subarachnoid space, where CSF is produced and flows, have been shown to drain Aβ^43,44^. But this role in the pathogenesis of AD is still disputed^36^. To date most research has focused on dorsal dural lymphatics, which sit above the brain. However, additional sites of CSF egress exist, including the lymphatic regions near cranial nerves at skull foramina. Further research into the role of these pathways in AD is warranted.

One cranial nerve lymphatic pathway, the cribriform plate lymphatics, is of particular interest in AD for several reasons. The cribriform plate (CP) is a portion of the ethmoid bone, and it is where the olfactory bulb (OB) sends olfactory nerve (ON) bundles to populate the nasal mucosa. As the olfactory nerve bundles pass through the cribriform plate, there are disruptions in the perineural subarachnoid layer allowing for direct sampling of the CSF by cribriform plate lymphatics^31,41,45^. Drainage at the cribriform plate lymphatics has been implicated to affect disease pathogenesis in autoimmune and ischemic stroke models of neuroinflammatory disease^31,46,47^, but have not been widely studied in settings of chronic neurodegeneration. In contrast to dorsal dural lymphatics, basal sites of CSF efflux including the CP lymphatics may have a larger capacity for CSF drainage due to their integration with nasal and nasopharyngeal lymphatics which have direct routes to cervical lymph nodes^48^. Moreover, clinical evidence using tracers for PET and MRI scan on AD patients and healthy volunteers have shown that AD patients have slower efflux of tracers at the nasal turbinates^49–52^. Specifically, they found that efflux of the ^18^F-THK5117 tracer had 23% lower ventricular clearance in AD patients and 66% fewer nasal egress sites in AD patients^50,51^, suggesting that while CSF efflux may be decreased within the brain parenchyma, it is further exacerbated in sites of CNS-associated lymphatic drainage. Furthermore, increased olfactory dysfunction in AD patients has been correlated as one of the five-year predictors of mortality in AD patients^53^, but the mechanism of olfactory dysfunction is still unclear.

To investigate the pathogenesis of AD, there are several established mouse models that we utilized for this research. APP/PS1 mice^54^ express a chimeric mouse/human amyloid precursor protein with the Swedish mutations, as well as a mutant human presenilin 1 (exon 9 deletion) in neurons, resulting in robust Aβ deposition. 5XFAD mice^55^ express mutant human APP with three mutations found in EOAD, as well as human PS1 harboring two EOAD mutations. 5XFAD mice are another model of Aβ deposition. These models have been extensively used in AD research.

In this paper, we demonstrate that these mouse models of AD recapitulate clinical findings of decreased CSF efflux at the cribriform plate. We demonstrate, for the first time, that the cribriform plate lymphatics are a site of Aβ deposition. This deposition of Aβ in APP/PS1 mice is associated with increased immune cell accumulation and decreased lymphatic vessel diameter at the cribriform plate, when compared to control mice. Additionally, an increase in cleaved caspase-3+ in lymphatic vessels of APP/PS1 mice suggests that Aβ results in cellular toxicity leading to apoptosis of cribriform plate lymphatic endothelial cells. These findings lay the groundwork for potential therapeutics to help modulate lymphatic vessel flow and mitigate lymphatic vessel cell death at skull foramina sites in AD.

## Results

### Decreased CSF efflux in APP/PS1 mice

PET and MRI tracer studies indicate that AD patients have decreased CSF efflux^9,50,51^. To understand if AD transgenic mouse models recapitulate those findings of reduced drainage at the cribriform plate, we utilized SPECT imaging in 24-month old aged APP/PS1 and age-matched control mice, to study the efflux rate of Technetium-99m (99mTc) at the cribriform plate and nasal turbinate area. The schematic of the experiment set-up is shown (**Figure 1A**). Briefly, mice received an intrathecal injection of 99mTc into the L5/L6 intervertebral space, to directly inject into the CSF-filled spinal subarachnoid space. After receiving the injection, mice were immediately imaged in the SPECT/CT scanner. Scans were taken every five minutes for a total of 35 minutes. The results show a significant decrease in 99mTc efflux at 5-minutes in APP/PS1 mice when compared to age-matched controls (**Figure 1B**), recapitulating clinical findings of decreased CSF-efflux in AD patients. We were able to gate and examine efflux at various anatomical sites along the cribriform plate efflux pathway (**Figure 1C-F**), demonstrating the presence of 99mTc at the nasal region in wildtype mice (**Figure 1C**). When compared in the same area at the same time-point in 24-month APP/PS1 mice, there was little contrast detected (**Figure 1D**). Interestingly, when we gated in the nasal lymphatic area, which has recently been shown to drain through the nasopharyngeal lymphatic plexus and directly into cervical lymph nodes, we were able to detect the same decrease in efflux as the cribriform plate (**Figure 1E-F**) demonstrating decrease CSF-efflux throughout the nasal drainage pathway. Due to size differences, proteins and Aβ oligomers within the CSF may behave differently during drainage. To better understand larger sized particulate drainage dynamics in APP/PS1 mice, we performed a 5 µL intracranial injection of FluoSpheres^TM^ with a size of 0.2 µm. 45 minutes after the bead injection, the mice were euthanized and perfused and tissue was collected (**Figure 1G**). We performed immunofluorescence analysis on the decalcified whole- head sections, and in aged-matched control mice observed bead collection between the OBs, in the area where cribriform plate lymphatics are located (**Figure 1H**, **1I**). This was not observed in 24-month old APP/PS1 mice (**Figure 1J**). The technical replicate quantification of these two images showed a trend of decreased beads present at the cribriform plate in APP/PS1 mice (**Figure 1K**) supporting the trend of decreased CSF drainage at the cribriform plate as seen in the SPECT/CT data. Overall, these results suggest that in APP/PS1 mice there is decreased CSF efflux through the cribriform plate that is not just due to age-related decline in lymphatic function, rather it is the consequence of APP/PS1 driven Aβ disease pathology.

**Figure 1.**
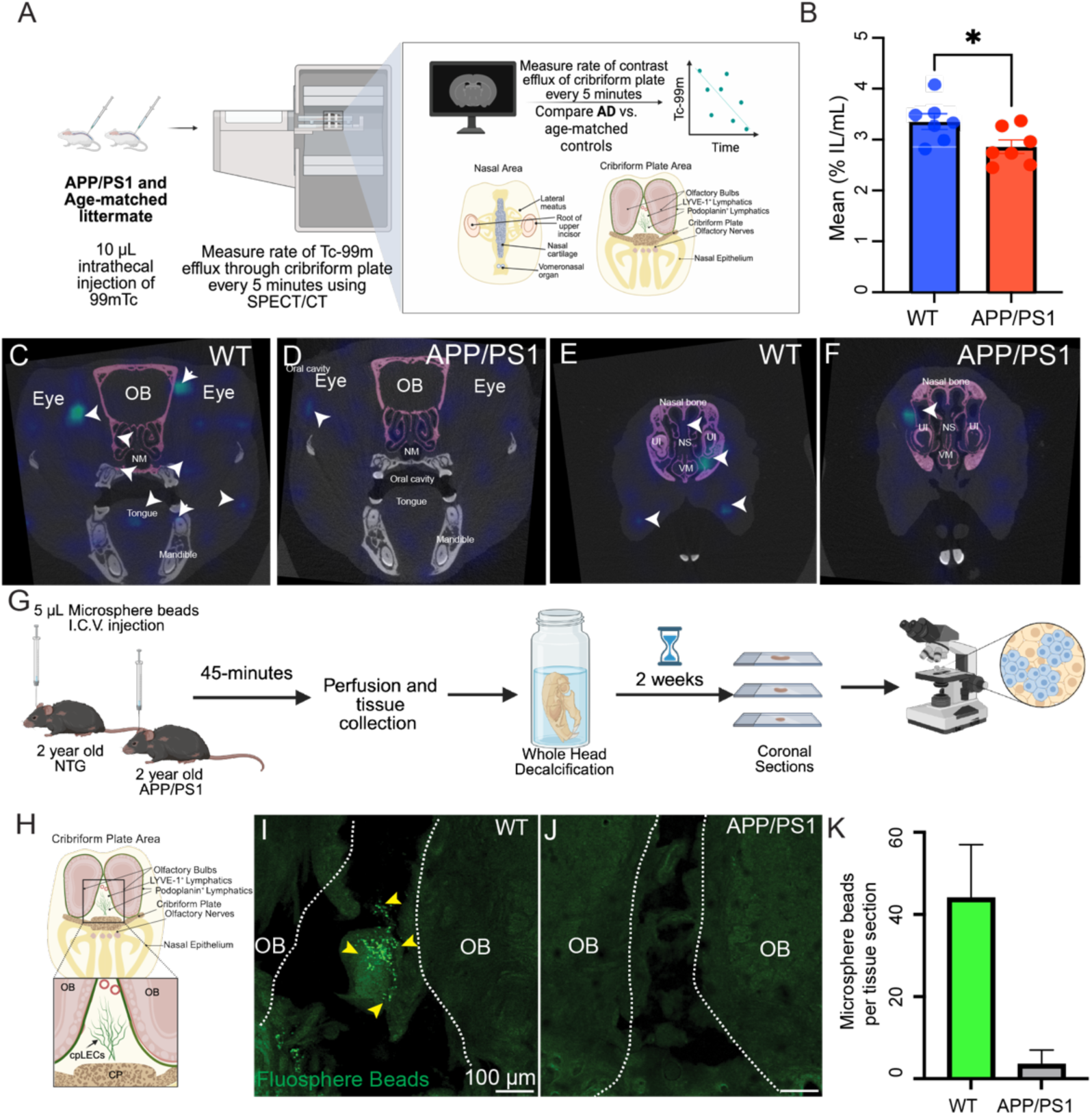
Decreased cerebrospinal fluid drainage in 24-month-old APP/PS1 mice. Schematic of SPECT/CT imaging to measure cerebrospinal fluid drainage through the cribriform plate (A). Briefly, 24-month-old control and 24-month-old APP/PS1 mice received a 10 µL intrathecal injection of 99m-technetium (99mTc) followed by SPECT scans every 5 minutes for 45 minutes. Graph of efflux at the cribriform plate comparing drainage between WT and APP/PS1 mice at 5 minutes post-injection; results of Student’s t-test p=0.033 (B). CT scans overlayed with SPECT scan at 5 minutes post injection highlighting 99mTc efflux through the cribriform plate in a 24-month-old wildtype (C) and APP/PS1 (D) mice. White arrows indicate the presence of 99mTc. Pink-shell defines the cribriform plate area. E-F) CT scan at 5-minutes in the nasal area of age-matched controls (E) and APP/PS1 mice (F). G) Schematic of bead injection experiment used to evaluate bead-drainage. Briefly, age-matched controls and 24- month-old APP/PS1 mice received a single intracerebral injection of 0.2 µm FluoSpheres^TM^ Blue (5 µL) into the right frontal lobe. 45-minutes later, mice were perfused, and tissues were collected for decalcification. H) Anatomical schematic of cribriform plate area highlighting cribriform plate lymphatics. G) Representative image of 24-month-old control mice with yellow arrows highlighting beads. H) Representative image of 24-month of APP/PS1 mice. I) Quantification of images from age-matched control and APP/PS1 mice. OB = olfactory bulbs, NS = nasal septum, UI = upper incisor, ON = olfactory nerves, and VM = vomeronasal organ.

### Aβ deposits in perineural space of olfactory nerve bundles

In mice, CSF and its contents have been demonstrated to accumulate and drain along the length of olfactory nerve bundles^35^. To investigate if Aβ is also present along CSF-interfacing olfactory nerve bundles (ON) at the cribriform plate in APP/PS1 mice, we performed immunostaining analysis on whole-head decalcified sections using specific antibodies against Aβ (D54D2), LYVE-1 (ALY7), and CD45 (30-F11). In 24-month old APP/PS1 mice, we detected strong Aβ aggregation within the OBs, meninges, and around ONs projecting through the cribriform plate skull foramina (**Figure 2C-D**).

**Figure 2.**
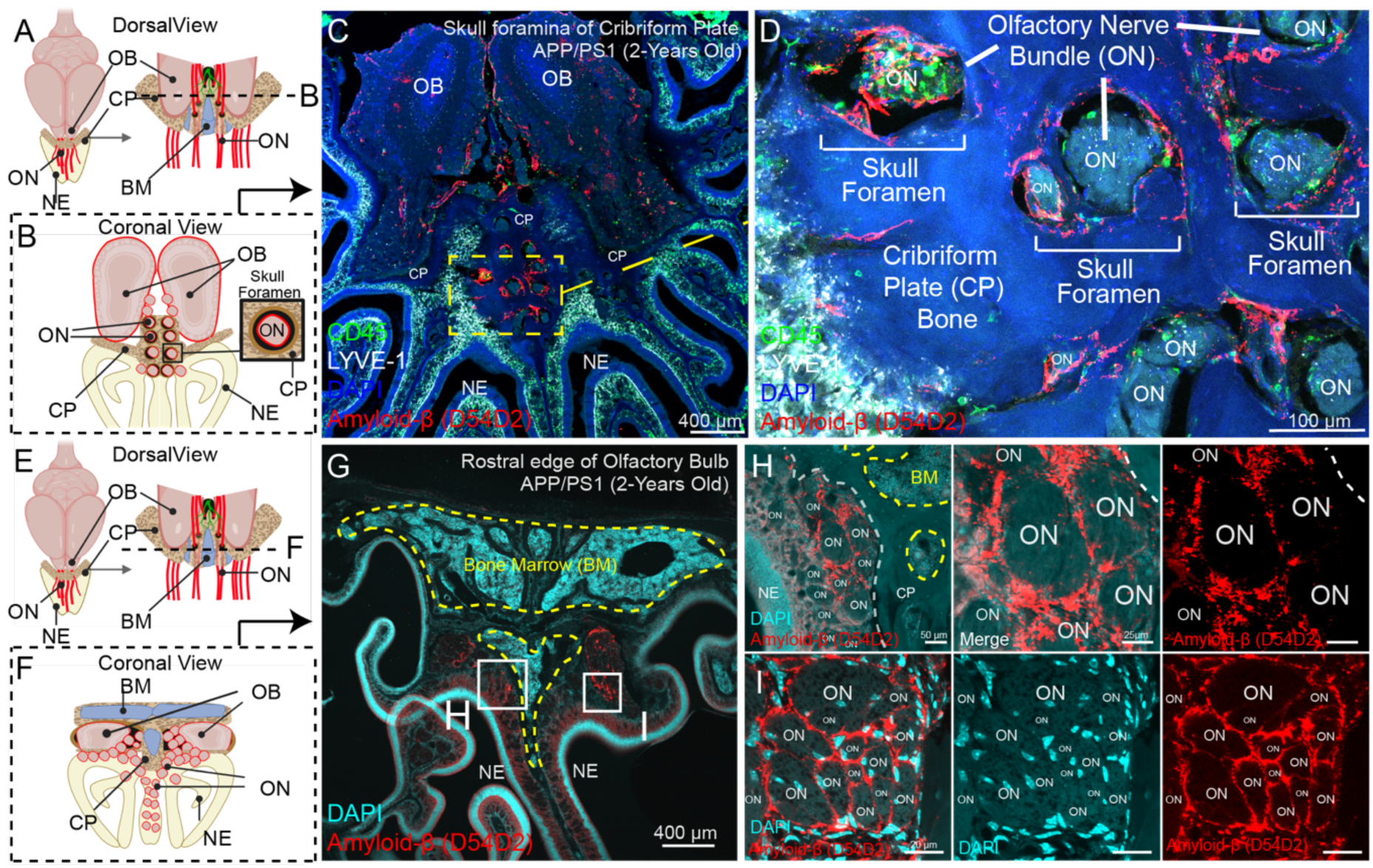
Aβ accumulates in the cribriform plate (CP) skull foramina and in the perineural spaces of olfactory nerve bundles in 24-month-old APP/PS1 mice. Anatomical scheme of the olfactory bulb (OB) - cribriform plate (CP) region from the dorsal view. Note the outward projecting olfactory nerve bundles (ON) through the CP into the nasal epithelium (NE) (A). Position “B” (dotted line) in the coronal view displays clear skull foramen where circular cross sectioned ONs transverse CP (B). Whole-head decalcified image (C) of CP foramina region of 24-month APP/PS1 mouse stained with for LYVE-1 (ALY7), CD45 (30-F11), Amyloid-beta (D54D2), and DAPI with zoomed in image (D) which shows Aβ in CP foramina and perineural space of ON. Anatomical scheme (E) where position “**F**” (dotted line) in the coronal view displays the rostral edge of OB and ON which have transversed the underside CP (F). Whole-head decalcified image of rostral CP region of 24-month-old APP/PS1 mouse (G) stained with for LYVE- 1 (ALY7), CD45 (30-F11), Amyloid-beta (D54D2), and DAPI with zoomed in images (H-I) which show Aβ in perineural space of ON. OB = olfactory bulbs, ON = olfactory nerves, NE = Nasal Epithelium, CP = Cribriform Plate, and BM = Bone Marrow.

Staining revealed “ring-like” patterns of Aβ accumulation in circular cross sections of ONs and inside skull foramen (**Figure 2C-D**). Additionally, in more rostral locations, near the front edge of the olfactory bulb, Aβ was observed extracranially accumulating beneath the cribriform plate along ONs in the lamina propria of nasal cavity (**Figure 2G**), again forming “ring-like” perineural Aβ aggregation patterns (**Figure 2H-I**).

Together this data supports that Aβ accumulates along CSF-interfacing olfactory nerve bundles.

### Aβ deposits within the cribriform plate lymphatic vessels

Cribriform lymphatic vessels directly associate and access the CSF-filled space along olfactory nerve bundles through gaps in E-cadherin^40,46^. To understand how AD disease progression is impacting cribriform plate lymphatic drainage, we performed immunofluorescence analysis on whole-head decalcified sections of 24-month old APP/PS1 mice and aged-control mice, staining for Aβ and lymphatics (LYVE-1). We show that LYVE-1 stains the cpLEC area between the OBs in both aged-control mice (**Figure 3A**) and APP/PS1 mice (**Figure 3B**). Most importantly, we show that Aβ is in close association with the LYVE-1+ cpLECs in APP/PS1 mice (**Figure 3B**). The staining for Aβ was specific as we do not see positive staining for Aβ in the aged-wild type mice (**Figure 3A**). We further confirmed that Aβ deposits in the cpLEC area using a different monoclonal antibody, clone 4G8, which is reactive to residues 17-24 of Aβ. When compared to the staining using the monoclonal antibody D54D2, which binds to residues near the N-terminus, a similar pattern of staining within the cpLEC area was observed (**Figure 3D**). Clinically, Aβ can be confirmed using antibody labeling, but it can also be detected through stains for the β-pleated sheet structure of Aβ. We performed paraffin-embedding on a decalcified 9-month old 5XFAD head, sectioned, and performed Congo Red staining. We imaged the sections using polarized light microscopy. The Congo Red staining additionally showed Aβ deposits within the olfactory bulb region (**Figure 3E-F**).

**Figure 3.**
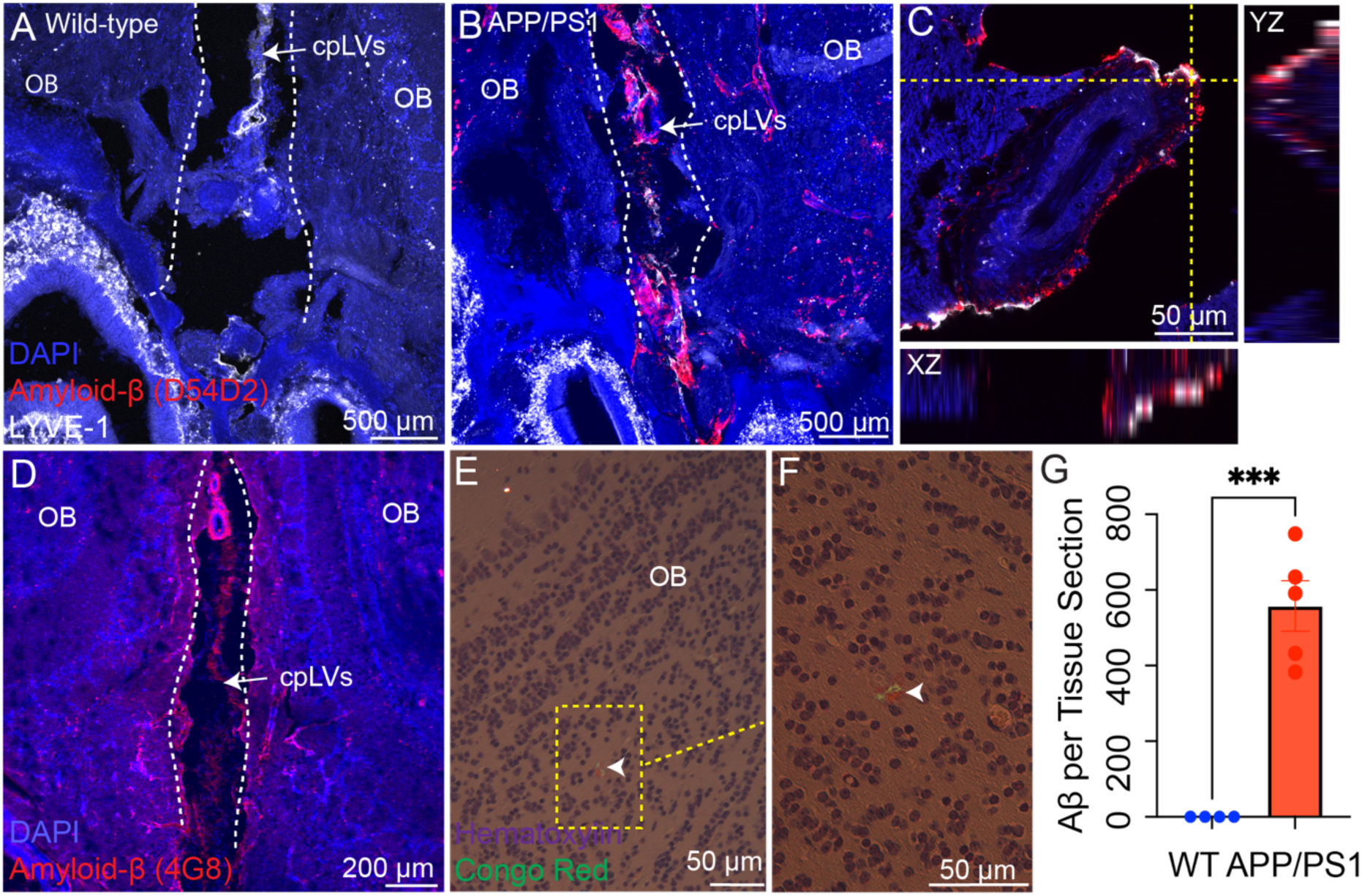
Aβ deposits in the cribriform plate lymphatics in AD transgenic mice. Healthy 24-month-old control (A) and 24-month-old APP/PS1 mice (B) were perfused and tissues were collected for whole-head decalcification. Sections were stained for amyloid-β (D54D2) and LYVE-1 (ALY7). 60X image showing close association of amyloid-β within the lymphatic vessels (C). Decalcified whole-head section of 24-month APP/PS1 mice were stained for DAPI and amyloid-β (4G8) (D). Decalcified whole-head section of 7-month-old 5XFAD was stained with Congo Red and Hematoxylin and imaged under polarized light microscopy (E) with magnified images shown (F). Quantification of amyloid-β in lymphatic area (G). OB = olfactory bulbs. Representative image of n=4-5 mice. Results of a Student’s unpaired t-test, p = 0.002.

To understand how much Aβ was depositing at the cribriform plate, we used Imaris software to analyze Aβ plaque counts. We counted plaques on the whole tissue section, which included the olfactory bulb (OB) and cpLEC area. We showed that there is a substantial amount of Aβ plaques present in the cpLEC and OB area (**Figure 3G**).

Taken together, we were able to verify and show that Aβ is depositing in close association with cpLECs in 24-month APP/PS1 mice.

### Aβ deposition is associated with lymphatic atrophy and increased immune cell recruitment

Our lab has previously shown that cpLECs undergo lymphangiogenesis as a result of neuroinflammation in the setting of conditions such as autoimmune encephalomyelitis (EAE) and transient middle cerebral artery occlusion (tMCAO)^31,46,47^. Since neuroinflammation is also observed in AD, we hypothesized that the close association of Aβ might impact the lymphatic network at the cribriform plate. Utilizing serial sectioning, we performed immunostaining analysis with antibodies specific for LYVE-1, a marker of cpLECs, and CD45, a common myeloid marker, to analyze immune cell recruitment. Representative images of the cpLEC area in 24-month old age-matched control and APP/PS1 mice are shown in Figure 4. We observed CD45+ cells in the LYVE-1 cpLEC area and in close association with Aβ aggregates (**Figure 4C-D**).

**Figure 4.**
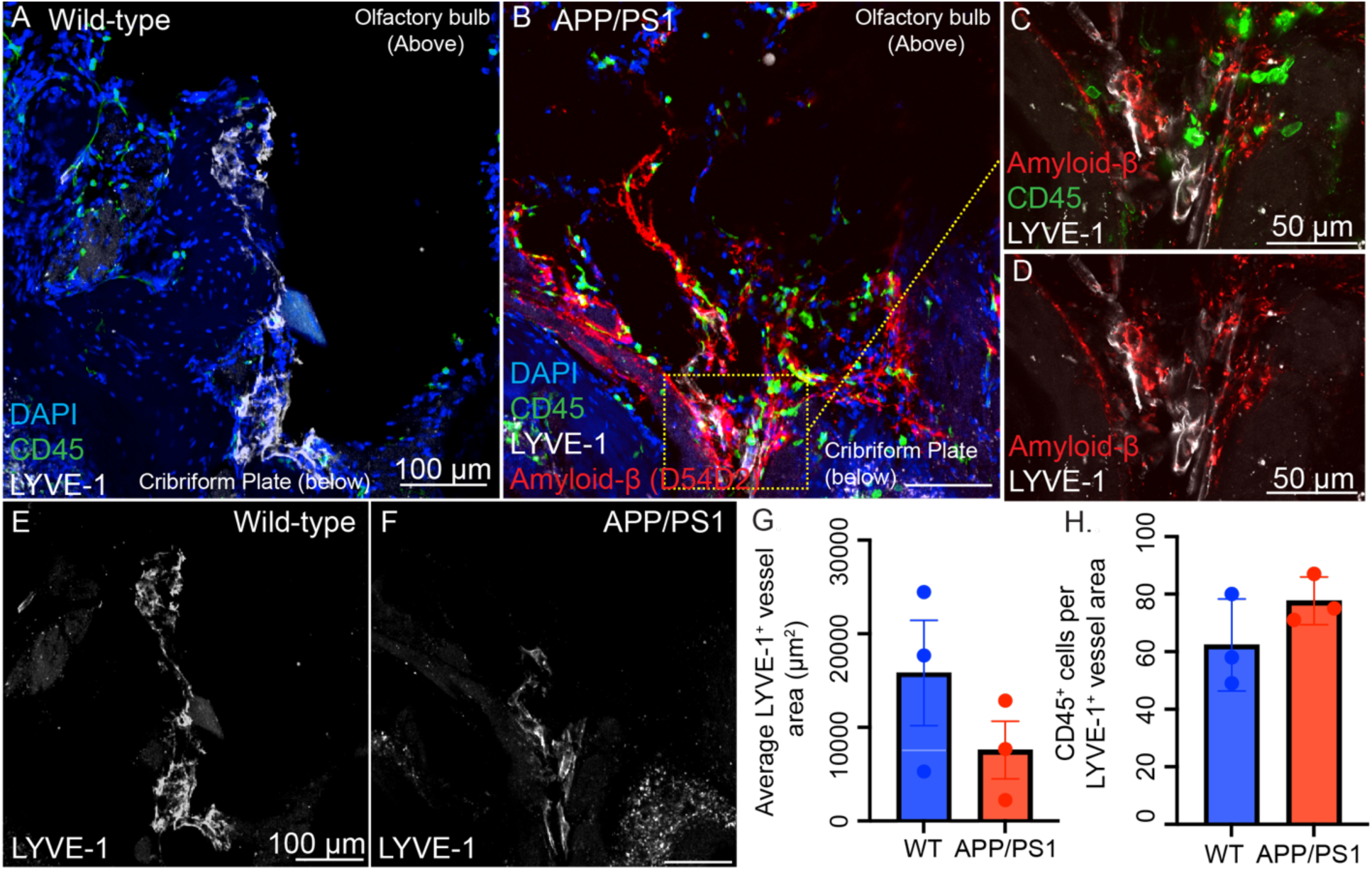
Cribriform plate lymphatics show a trend of decreased vessel area and increased CD45+ immune cell recruitment in APP/PS1 mice. Healthy 24-month-old control (A) and 24-month-old APP/PS1 (B) were euthanized and perfused prior to tissue collection and whole-head decalcification. Sections were stained for CD45 (30-F11), LYVE-1 (ALY7), DAPI (A) and Amyloid-β (D54D2) (B). Several areas of CD45+ cells accumulating near LYVE-1+ vessels were shown (C). Close association of Amyloid-β and LYVE-1 was seen (D). Lymphatic vessel area and structure is altered in APP/PS1 mice (F) when compared to healthy (E) mice. LYVE-1+ vessel area and CD45+ cells of the shown images (A and B) were analyzed on ImageJ and results are shown (G and H). Three serial sections were taken of healthy and APP/PS1 mice and LYVE-1+ vessel area was measured. CD45+ cells at the LYVE-1+ vessel area (as shown in E and F) were counted in the three serial sections using Cell Counter on ImageJ. A Student’s unpaired t-test was performed on GraphPad Prism with p = 0.2684 (G) and p = 0.2139 (H). There is a trend of decreased LYVE-1 vessel area (G) and increased CD45+ immune cell recruitment (H).

Additionally, through visualization of LYVE-1+ area, we observed that the CP lymphatic area in the aged-match controls (**Figure 4E**) was larger than the APP/PS1+ mice (**Figure 4F**). We analyzed the LYVE-1+ area rostral and caudal to the representative images, and we show that there is a trend of decreased LYVE-1+ area (**Figure 4G**) in APP/PS1 mice. In addition, we performed the same analysis but counting the number of CD45+ cells present within the cpLEC area, excluding CD45+ cells that were in the OB or within the nasal epithelium below the CP (**Figure 4H**). This analysis showed a trend of increased immune cell recruitment. Overall, we had the important finding of decreased LYVE-1+ vessel area in APP/PS1 mice in comparison to age matched controls, as well as trending increased CD45+ immune cell recruitment to the cpLEC area. Future work will delineate what type of immune cells have been recruited to this area and are interacting with Aβ aggregates.

### Aβ deposition results in increased cell death of lymphatic vessels

The described decrease in LYVE-1+ vessel area potentially represents lymphatic atrophy. Therefore, we sought to determine what was driving this decrease in lymphatic vessel area. We hypothesized that inflammation and toxicity caused by Aβ deposits may be resulting in cell death of the cribriform plate lymphatic vessels. To assess for cell death, we stained for cleaved caspase-3 in decalcified whole-head sections of 24- month old APP/PS1 and age-matched controls. We focused our analysis at the area of the cpLECs between the olfactory bulbs. We showed that in 24-month old age-matched control mice there was minimal cleaved caspase-3 in podoplanin+ LECs (**Figure 5A**).

**Figure 5.**
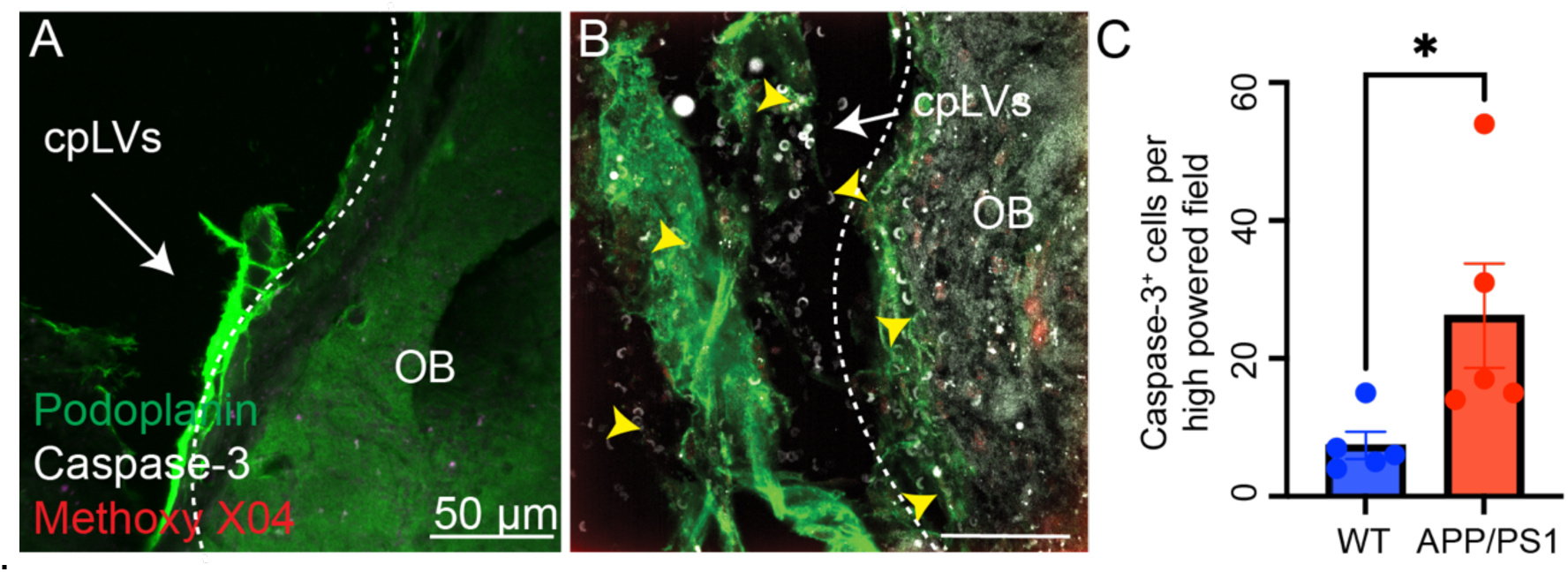
Increased cleaved caspase-3 staining podoplanin^+^ lymphatic area of APP/PS1 mice. Healthy 24-month-old control (A) and 24-month-old APP/PS1 (B) mice were euthanized and perfused; tissues were collected, and heads were decalcified. Images at 60x magnification of tissue sections stained with podoplanin (eBio8.1.1), Caspase-3 (Asp175), and Methoxy X04. Images were taken between the olfactory bulbs in the lymphatic area. Podoplanin areas on the tissue sections were randomly selected for imaging, and the caspase-3^+^ areas were counted using ImageJ analysis. A Student’s t-test was performed using GraphPad Prism and p-value = 0.0226. Arrows indicate areas of cleaved caspase-3 staining.

However, in 24-month old APP/PS1 mice, there was increased cleaved caspase-3 staining (**Figure 5B and 5C**). This suggests that the decrease in LEC vessel area is likely due to cell death induced by Aβ deposition.

## Discussion

Alzheimer’s disease is increasingly being recognized as a systemic disease with the pathology extending far beyond the brain itself. It is within these peripheral systems that we can elucidate disease pathogenesis and discover innovative therapeutic approaches. In AD disease pathogenesis, there is accumulation of Aβ and phosphorylated tau leading to neurotoxicity, neurodegeneration, and subsequent symptoms. The clearance and disposal of these harmful proteins are crucial to disease pathogenesis and represent an innovative opportunity for treatments. The waste-management system of the brain relies on parenchymal immune responses, glymphatic clearance, and a complex network of CNS-adjacent lymphatic vessels. In this study, we show that Aβ accumulates at the cribriform plate lymphatics, an important site of CSF efflux, coating lymphatic vessels. In addition, we show that Aβ deposition is associated with decreased CSF efflux, lymphatic vessel atrophy, and lymphatic vessel death.

In the context of AD, the dorsal meningeal lymphatics have been investigated, although with conflicting results^43,44,56^. It has been debated whether Aβ truly deposits within the meningeal lymphatic vessels, with one group showing there was positive Aβ staining within whole-mount dura meningeal lymphatic vessels, while lymphatic ablation worsened AD disease pathogenesis in mice treated with anti-Aβ passive immunotherapy^43,44^. Another group examining whole-mount dura meningeal lymphatic vessels observed that Aβ staining was actually within bridging veins in the meningeal layer and that lymphatic ablation did not worsen disease pathology^36^. There are several possible explanations for these discrepancies, including differences in the methods used to ablate the lymphatics and the potential compensation by other CSF efflux sites, which are areas of further research. Furthermore, additional studies have implicated the glymphatic system and cranial nerve roots in AD pathology; however, no such study has specifically focused on the cribriform plate lymphatics. Broadly, the cribriform plate has been shown to facilitate fluid and antigen drainage and plays a critical regulatory role in the setting of autoimmunity and stroke^31,46,46,47^. In this study, we describe the cribriform plate as a site of amyloid deposition in AD, warranting further studies on subsequent immune regulation and modulation of this anatomical niche.

One challenge within the field of CNS lymphatics has been the ability to quantify CSF flow along its egress pathways. By combining high-quality CT images with multiple single-photon emission computed tomography (SPECT) over a time-course, we observed that CSF efflux is decreased in AD transgenic models. By combining the data from 99mTc efflux with the CT images, we were able to delineate the anatomic site where efflux of the tracer was occurring. Previously, we have been limited by the resolution of the MRI imaging, but the combination of high-quality CT images with SPECT imaging provides excellent spatial resolution. Additionally, with this technique, we were able to verify areas of CSF flow that have been previously reported; specifically, we were able to visualize tracer efflux through the ocular system as well as the nasopharynx area^48,57^. Utilizing this method, we were able to see in 24-month APP/PS1 mice have decreased CSF efflux at 5-minutes through the cribriform plate area. As the efflux was decreased in AD in comparison to age-match controls, it suggests that the decrease in CSF efflux is due to AD-associated pathology rather than age. Specifically, it is an exacerbation of an already declining system, as it is established that with age, the function of CNS-associated lymphatics decline.^44,58–60^ It is possible that with the build-up of Aβ over time to a stressed system, there is eventually a point at which the body’s compensatory drainage abilities are lost, resulting in disease pathogenesis acceleration. We will need to examine other time-points to better determine this relationship. However, these results recapitulate clinical findings in AD patients, and it demonstrates that the cribriform plate is an important CSF efflux site in mice.

After discovering that CSF efflux was decreased in APP/PS1 mice, we assessed if the cribriform plate was an area of Aβ deposition and aggregation, hypothesizing that Aβ aggregates were acting as a plug at this important drainage site. We demonstrated that there is heavy deposition of Aβ at the cribriform plate, showing that Aβ lines the olfactory nerves as they project through the cribriform plate, which is never been demonstrated before. In addition to lining the olfactory nerves, there is heavy deposition of Aβ, verified through multiple methods, in close association with the cribriform plate lymphatic endothelial cells. It has been debated if Aβ truly deposits within CNS lymphatics with contradictory results in the dorsal meningeal structures^43,44,56^ and these results show for the first-time that Aβ does deposit within lymphatic structures at the cribriform plate lymphatics.

Our lab and other groups have demonstrated that the cribriform plate lymphatics undergo lymphangiogenesis in response to neuroinflammation. Despite neuroinflammation being a recognized hallmark of AD disease pathology, the cribriform plate lymphatics did not undergo lymphangiogenesis in response to disease progression at the 24-month time point. Instead, unexpectedly, the lymphatic vessels underwent atrophy. Moreover, there was increased immune cell recruitment in APP/PS1 mice, suggesting that there is an inflammatory process occurring or potentially, the immune cells efflux is also hampered by the decreased CSF efflux seen during the pathology. This led us to explore the possibility of the lymphatic vessels undergoing apoptosis. There is evidence in other CNS-associated lymphatics of Aβ disrupting function. Deposition of Aβ has been described along the optic nerve resulting in absence and loss of polarity of aquaporin (AQP)-4 channels, which play an important role in CSF fluid flow through the glymphatic system^57^. Additionally, it has been well established that Aβ can deposit within blood vessels, resulting in cerebral amyloid angiopathy, which can result in endothelial cell death and damage^61^. However, it has not been shown before to result in cell death of lymphatic endothelial cells (LECs).

Moreover, caspase-3 has been implicated in the processing of amyloid precursor protein with higher levels of caspase-3 positively corresponding with AD pathology as well as high levels of caspase-3 in close association with neurofibrillary tangles potentially playing a role in tau truncation^62–64^. We hypothesize that this lymphatic atrophy potentially represents the end of the disease spectrum for AD. It is possible that in early stages of disease there is lymphangiogenesis to promote Aβ efflux, but over time the accumulation of Aβ becomes too high, resulting in decreased CSF flow and eventual cell death of the cribriform plate lymphatic endothelial cells. Future studies will aim to examine this process at earlier time points as well as delineate which specific immune cells are being recruited to the cpLEC area.

Overall, we demonstrate that, in aged-APP/PS1 mice, the cribriform plate lymphatics are an area impacted by AD disease pathogenesis. Utilizing multiple methods of SPECT/CT analysis and bead injection, we demonstrated that CSF efflux is decreased at the cribriform plate in APP/PS1 mice. We also demonstrated that Aβ deposits at the cribriform plate in close association with cribriform plate lymphatic endothelial cells.

Moreover, we show that at 24-months of age, there is a decrease in lymphatic vessel area and increased immune cell infiltration, which is suggestive of vessel atrophy.

Finally, we showed that cribriform plate lymphatic endothelial cells are undergoing cell death in association with Aβ. Together these findings suggest that skull foramina sites, particularly the CSF efflux routes along perineural lymphatics, are relevant locations to further investigate AD-related amyloid accumulation, immune response, and lymphatic dysfunction. These findings lay the groundwork for novel investigations along the brain’s borders to generate new therapies and biomarkers.

## Materials and Methods

### Animals

24-month-old male and female B6.Cg-Tg(APPswe,PSEN1dE9)85Dbo/Mmjax (APP/PS1)^54^ mice were obtained from were obtained from the Jackson Laboratory and maintained as hemizygous animals in a congenic C57BL/6J background. Male and female B6.Cg-Tg(APPSwFlLon,PSEN1*M146L*L286V)6799Vas/Mmjax (5XFAD)^55^ were a generous gift from the laboratory of Dr. Tyler Ulland. Mice were initially purchased from the Mutant Mouse Resource and Research Centers (MMRRC). 5XFAD were used at 9-months of age. All experiments were conducted in accordance with the guidelines from the National Institutes of Health and the University of Wisconsin-Madison Institutional Animal Care and Use Committee.

### SPECT/CT Imaging

SPECT/CT Imaging was performed on MILABs U-SPECT/CT^UHR^ (Ultra-high resolution) system. Animals were administered isoflurane through a nose cone to anesthetize for the intrathecal injection and scanning. Prior to imaging, APP/PS1 and age-matched control mice were given a 10 µL injection of 99m-technetium into the L5/L6 intervertebral space. After receiving the injection, mice were immediately imaged in the SPECT/CT scanner with scans focused on the cribriform plate. Images were taken every 5 minutes to understand efflux through the cribriform plate area. Mice were monitored throughout the scan, and they were monitored after the scan to ensure proper recovery. The scans were analyzed using Imalytics.

### Histology

To collect tissues, mice were anesthetized with excess isoflurane and following respiratory rate cessation, mice were transcardially perfused with PBS followed by 4% paraformaldehyde. Tissues were collected for histological analysis including the whole head following decapitation. The whole head was fixed in 4% paraformaldehyde for additional 24 hours, and then it was decalcified in 14% EDTA for two weeks. The EDTA was changed every two days. Following decalcification, the tissues were rehydrated in 40% sucrose for 48 hours. The tissues were embedded in Tissue-Tek OCT Compound, frozen on dry ice, and stored at -80°C. Tissues were sectioned on a Leica CM1800 cryostat at section thickness of 40 µm and mounted on Superfrost Plus microscope slides. The slides were stored at -80°C.

### Immunofluorescence and confocal microscopy

For immunofluorescence, slides were thawed and then washed with PBS with 1% bovine albumin serum (BSA) for 5 minutes. Blocking buffer was then added for one hour. Sections were stained with rabbit anti-amyloid-beta (1:100, D54D2, Cell Signaling Technology), rat anti-LYVE-1-eFluor-660 conjugated (1:200, ALY7, 50-044-82, Thermo Fisher Scientific), rat anti-Podoplanin-eFluor488 conjugated (1:500, 53-5381-80, eBioscience), rat anti-CD45-AlexaFluor488 (1:200, 30-F11, Invitrogen, 53-0451-82), and Methoxy X04 (1:1000). Incubation with the antibodies were performed in staining solution of 1% BSA and 0.1% Triton X overnight at 4°C. For sections stained with rabbit anti-amyloid-beta 17-24 (1:200, 4G8, BioLegend 800701) and cleaved caspase-3 (1:100, 9662, Cell Signaling Technology), sections were incubated in staining solution of 5% BSA, 5% donkey serum, and 0.3% Triton X overnight at 4°C. Following staining with primary antibody solutions, sections were washed three times for ten minutes in blocking buffer, and slides were incubated with the appropriate secondary antibody staining solution for two hours at room temperature. Secondary antibodies used were donkey anti-rabbit AlexaFluor568 (1:500, Invitrogen, A10042) and donkey anti-rabbit AlexaFluor647 (1:500, Invitrogen, A31573). Following secondary staining, sections were again washed three times for ten minutes and mounted with Prolong Gold mounting medium with DAPI.

Images were acquired using an Olympus Fluoview FV1200 confocal microscope and a Nikon A1R Confocal Microscope. Laser imaging settings were kept the same across the replicates. Images were analyzed using FIJI software utilizing the same brightness and contrast levels.

### Congo Red Staining and polarized microscopy

Congo red staining was performed on decalcified, whole-head sections embedded in paraffin. Sections were deparaffinized, stained with Harris hematoxylin, rinsed with 95% ethanol, stained with Alkaline Congo Red Solution for 20 minutes, rinsed with NaCl ethanol followed by absolute alcohol twice, cleared with xylene, and mounted in synthetic mounting media. Images were acquired using a Nikon Eclipse E600 microscope fitted with polarized light filters and a Nikon DSFi2 camera. Images were taken at 20X resolution.

### Statistical analysis

Statistical analysis was performed using GraphPad Prism. Where indicated, a unpaired Student’s t-test was performed.

## Acknowledgements

I would like to thank members of our laboratory for helpful discussions and constructive critiques on this project including Melinda Herbath, Jenna Port, Thiunuwan Priyathilaka Thanthrige, Kristof Kovacs, and Laura Schmitt Brunold. Thank you to fantastic undergraduate research assistants, Dhruv Bansal, Abigail M. Sehmer, Athena N. Kafkas, Mohan Kumar, and Simon Federico Woen Ordonez.

The authors would like to acknowledge the Cancer Center Support Grant: NCI P30 CA014520, University of Wisconsin Small Animal Imaging & Radiotherapy Facility and NIH S10OD028670-01 for supporting this work. Additionally, we thank the University of Wisconsin Translational Research Initiatives in Pathology (TRIP) Laboratory, supported by UW Department of Pathology and Laboratory Medicine, UWCCC (P30CA14520) and the Office of the Director NIH (S10 OD023526). We thank the Optical Imaging Core, supported by 1S10034394-01.

The work was supported by NIH grant 5R01NS126595 to Z.F. Training support to S.M.V. through 5T32GM135119 and 5T32GM140935. The authors have no conflict of interest to declare.

## References

1. 2024 Alzheimer’s disease facts and figures. Alzheimers Dement. J. Alzheimers Assoc. 20, 3708–3821 (2024).

2. Skaria, A. P. The economic and societal burden of Alzheimer disease: managed care considerations. Am. J. Manag. Care 28, S188–S196 (2022).

3. Cummings, J., Lee, G., Ritter, A., Sabbagh, M. & Zhong, K. Alzheimer’s disease drug development pipeline: 2019. Alzheimers Dement. N. Y. N 5, 272–293 (2019).

4. Yiannopoulou, K. G. & Papageorgiou, S. G. Current and Future Treatments in Alzheimer Disease: An Update. J. Cent. Nerv. Syst. Dis. 12, 1179573520907397 (2020).

5. Yaghmaei, E. et al. Combined use of Donepezil and Memantine increases the probability of five-year survival of Alzheimer’s disease patients. Commun. Med. 4, 1– 8 (2024).

6. Mintun, M. A. et al. Donanemab in Early Alzheimer’s Disease. N. Engl. J. Med. 384, 1691–1704 (2021).

7. Dyck, C. H. van et al. Lecanemab in Early Alzheimer’s Disease. N. Engl. J. Med. 388, 9–21 (2023).

8. Graeber, M. B. et al. Rediscovery of the case described by Alois Alzheimer in 1911: historical, histological and molecular genetic analysis. Neurogenetics 1, 73–80 (1997).

9. Hardy, J. & Higgins, G. A. Alzheimer’s disease: the amyloid cascade hypothesis. Science 256, 184–5 (1992).

10. Hardy, J. & Selkoe, D. J. The amyloid hypothesis of Alzheimer’s disease: progress and problems on the road to therapeutics. Science 297, 353–356 (2002).

11. Selkoe, D. J. & Hardy, J. The amyloid hypothesis of Alzheimer’s disease at 25 years. EMBO Mol. Med. 8, 595–608 (2016).

12. Selkoe, D. J. et al. Beta-amyloid precursor protein of Alzheimer disease occurs as 110- to 135-kilodalton membrane-associated proteins in neural and nonneural tissues. Proc. Natl. Acad. Sci. U. S. A. 85, 7341–7345 (1988).

13. Berezovska, O. et al. Familial Alzheimer’s disease presenilin 1 mutations cause alterations in the conformation of presenilin and interactions with amyloid precursor protein. J. Neurosci. Off. J. Soc. Neurosci. 25, 3009–3017 (2005).

14. Levitan, D. et al. PS1 N- and C-terminal fragments form a complex that functions in APP processing and Notch signaling. Proc. Natl. Acad. Sci. U. S. A. 98, 12186– 12190 (2001).

15. Mawuenyega, K. G. et al. Decreased Clearance of CNS Amyloid-β in Alzheimer’s Disease. Science 330, 1774 (2010).

16. Cacace, R., Sleegers, K. & Van Broeckhoven, C. Molecular genetics of early- onset Alzheimer’s disease revisited. Alzheimers Dement. 12, 733–748 (2016).

17. Carmona, S., Hardy, J. & Guerreiro, R. Chapter 26 - The genetic landscape of Alzheimer disease. in Handbook of Clinical Neurology (eds. Geschwind, D. H., Paulson, H. L. & Klein, C.) vol. 148 395–408 (Elsevier, 2018).

18. Chen, Y., Strickland, M. R., Soranno, A. & Holtzman, D. M. Apolipoprotein E: Structural Insights and links to Alzheimer’s Disease Pathogenesis. Neuron 109, 205– 221 (2021).

19. Kim, J., Basak, J. M. & Holtzman, D. M. The Role of Apolipoprotein E in Alzheimer’s Disease. Neuron 63, 287–303 (2009).

20. Mahley, R. W., Weisgraber, K. H. & Huang, Y. Apolipoprotein E: structure determines function, from atherosclerosis to Alzheimer’s disease to AIDS. J. Lipid Res. 50 **Suppl**, S183–188 (2009).

21. Corder, E. H. et al. Gene Dose of Apolipoprotein E Type 4 Allele and the Risk of Alzheimer’s Disease in Late Onset Families. Science 261, 921–923 (1993).

22. Raulin, A.-C. et al. ApoE in Alzheimer’s disease: pathophysiology and therapeutic strategies. Mol. Neurodegener. 17, 72 (2022).

23. Bloom, G. S. Amyloid-β and Tau: The Trigger and Bullet in Alzheimer Disease Pathogenesis. JAMA Neurol. 71, 505–508 (2014).

24. Noble, W., Hanger, D. P., Miller, C. C. J. & Lovestone, S. The Importance of Tau Phosphorylation for Neurodegenerative Diseases. Front. Neurol. 4, 83 (2013).

25. Takashima, A., Noguchi, K., Sato, K., Hoshino, T. & Imahori, K. Tau protein kinase I is essential for amyloid beta-protein-induced neurotoxicity. Proc. Natl. Acad. Sci. U. S. A. 90, 7789–7793 (1993).

26. Heneka, M. T. et al. Neuroinflammation in Alzheimer disease. Nat. Rev. Immunol. 25, 321–352 (2025).

27. Hansen, D. V., Hanson, J. E. & Sheng, M. Microglia in Alzheimer’s disease. J. Cell Biol. 217, 459–472 (2018).

28. Zhou, Y. et al. Human and mouse single-nucleus transcriptomics reveal TREM2- dependent and TREM2-independent cellular responses in Alzheimer’s disease. Nat. Med. 26, 131–142 (2020).

29. Wang, Y. et al. TREM2-mediated early microglial response limits diffusion and toxicity of amyloid plaques. J. Exp. Med. 213, 667–675 (2016).

30. Keren-Shaul, H. et al. A Unique Microglia Type Associated with Restricting Development of Alzheimer’s Disease. Cell 169, 1276–1290.e17 (2017).

31. Hsu, M., Laaker, C., Sandor, M. & Fabry, Z. Neuroinflammation-Driven Lymphangiogenesis in CNS Diseases. Front. Cell. Neurosci. 15, (2021).

32. Louveau, A. et al. Structural and functional features of central nervous system lymphatic vessels. Nature 523, 337–341 (2015).

33. Engelhardt, B. & Ransohoff, R. M. The ins and outs of T-lymphocyte trafficking to the CNS: anatomical sites and molecular mechanisms. Trends Immunol. 26, 485–495 (2005).

34. Alitalo, K. The lymphatic vasculature in disease. Nat. Med. 17, 1371–1380 (2011).

35. Proulx, S. T. Cerebrospinal fluid outflow: a review of the historical and contemporary evidence for arachnoid villi, perineural routes, and dural lymphatics. Cell. Mol. Life Sci. 78, 2429–2457 (2021).

36. Antila, S. et al. Development and plasticity of meningeal lymphatic vessels. J. Exp. Med. 214, 3645–3667 (2017).

37. Kipnis, J. The anatomy of brainwashing. Science 385, 368–370 (2024).

38. Rasmussen, M. K., Mestre, H. & Nedergaard, M. The glymphatic pathway in neurological disorders. Lancet Neurol. 17, 1016–1024 (2018).

39. Mestre, H., Mori, Y. & Nedergaard, M. The Brain’s Glymphatic System: Current Controversies. Trends Neurosci. 43, 458–466 (2020).

40. Spera, I. et al. Open pathways for cerebrospinal fluid outflow at the cribriform plate along the olfactory nerves. EBioMedicine 91, 104558 (2023).

41. Madarasz, A., Xin, L. & Proulx, S. T. Clearance of erythrocytes from the subarachnoid space through cribriform plate lymphatics in female mice. EBioMedicine 107, 105295 (2024).

42. Kaag Rasmussen, M., et al. Trigeminal ganglion neurons are directly activated by influx of CSF solutes in a migraine model. Science 385, 80–86 (2024).

43. Da Mesquita, S. et al. Meningeal lymphatics affect microglia responses and anti- Aβ immunotherapy. Nature 593, 255–260 (2021).

44. Da Mesquita, S. et al. Functional aspects of meningeal lymphatics in ageing and Alzheimer’s disease. Nature 560, 185–191 (2018).

45. Ma, Q. et al. Rapid lymphatic efflux limits cerebrospinal fluid flow to the brain. Acta Neuropathol. (Berl.) 137, 151–165 (2019).

46. Hsu, M. et al. Neuroinflammation creates an immune regulatory niche at the meningeal lymphatic vasculature near the cribriform plate. Nat. Immunol. 23, 581– 593 (2022).

47. Choi, Y. H. et al. Dual role of vascular endothelial growth factor-C in post-stroke recovery. J. Exp. Med. 222, e20231816 (2025).

48. Yoon, J.-H. et al. Nasopharyngeal lymphatic plexus is a hub for cerebrospinal fluid drainage. Nature 625, 768–777 (2024).

49. Blennow, K., de Leon, M. J. & Zetterberg, H. Alzheimer’s disease. Lancet Lond. Engl. 368, 387–403 (2006).

50. Li, Y. et al. Decreased CSF clearance and increased brain amyloid in Alzheimer’s disease. Fluids Barriers CNS 19, 21 (2022).

51. de Leon, M. J. et al. Cerebrospinal Fluid Clearance in Alzheimer Disease Measured with Dynamic PET. J. Nucl. Med. Off. Publ. Soc. Nucl. Med. 58, 1471– 1476 (2017).

52. Tarasoff-Conway, J. M. et al. Clearance systems in the brain-implications for Alzheimer disease. Nat. Rev. Neurol. 11, 457–470 (2015).

53. Schubert, C. R. et al. Olfaction and the 5-year incidence of cognitive impairment in an epidemiological study of older adults. J. Am. Geriatr. Soc. 56, 1517–1521 (2008).

54. Jankowsky, J. L. et al. Mutant presenilins specifically elevate the levels of the 42 residue beta-amyloid peptide in vivo: evidence for augmentation of a 42-specific gamma secretase. Hum. Mol. Genet. 13, 159–170 (2004).

55. Oakley, H. et al. Intraneuronal β-Amyloid Aggregates, Neurodegeneration, and Neuron Loss in Transgenic Mice with Five Familial Alzheimer’s Disease Mutations: Potential Factors in Amyloid Plaque Formation. J. Neurosci. 26, 10129–10140 (2006).

56. Antila, S. et al. Sustained meningeal lymphatic vessel atrophy or expansion does not alter Alzheimer’s disease-related amyloid pathology. *Nat*. Cardiovasc. Res. 3, 474–491 (2024).

57. Cao, Q. et al. Transport of β-amyloid from brain to eye causes retinal degeneration in Alzheimer’s disease. J. Exp. Med. 221, e20240386 (2024).

58. Rego, S., Sanchez, G. & Da Mesquita, S. Current views on meningeal lymphatics and immunity in aging and Alzheimer’s disease. Mol. Neurodegener. 18, 55 (2023).

59. Jiang, H. et al. Overview of the meningeal lymphatic vessels in aging and central nervous system disorders. Cell Biosci. 12, 202 (2022).

60. Liu, Q. et al. Age-related changes in meningeal lymphatic function are closely associated with vascular endothelial growth factor-C expression. Brain Res. 1833, 148868 (2024).

61. Take, Y. et al. Amyloid β aggregation induces human brain microvascular endothelial cell death with abnormal actin organization. Biochem. Biophys. Rep. 29, 101189 (2022).

62. Gamblin, T. C. et al. Caspase cleavage of tau: linking amyloid and neurofibrillary tangles in Alzheimer’s disease. Proc. Natl. Acad. Sci. U. S. A. 100, 10032–10037 (2003).

63. Louneva, N. et al. Caspase-3 Is Enriched in Postsynaptic Densities and Increased in Alzheimer’s Disease. Am. J. Pathol. 173, 1488–1495 (2008).

64. Su, J. H., Zhao, M., Anderson, A. J., Srinivasan, A. & Cotman, C. W. Activated caspase-3 expression in Alzheimer’s and aged control brain: correlation with Alzheimer pathology. Brain Res. 898, 350–357 (2001).

